# Urine proteomic analysis of patients with schizophrenia

**DOI:** 10.1101/2023.11.11.566683

**Authors:** Chenyang Zhao, Fang Dong, Fuchun Zhou, Yuhang Huan, Jian Yang, Youhe Gao

## Abstract

We tried to explore the difference of urinary proteome between unmedicated schizophrenia patients and normal people through a small number of cases. The results showed that a total of 35 differential proteins were screened in the schizophrenia group compared with the healthy control group. Through random grouping evaluation, it has 91.4 % credibility. Fifteen of the differentially expressed proteins were reported to be related to schizophrenia mechanism, drug target or nervous system regulation. Among them, the aromatic amino acid decarboxylase related to the pathogenesis of schizophrenia can distinguish all 10 patients and 9 normal people with 100 % accuracy in this study, and the AUC value of 17 proteins is greater than or equal to 0.9. The biological pathways enriched by differentially expressed proteins include ephrin receptor signaling pathway, positive regulation of long-term potentiation (LTP), etc. This study shows that urine proteomics can reflect the difference between schizophrenia and healthy controls, and has the potential as a diagnostic marker.

## 1 Introduction

Schizophrenia, also known as mental disorders, integration disorders, is a group of unknown etiology of chronic diseases, involving sensory perception, thinking, emotion and behavior and other aspects of the obstacles and mental activity uncoordinated^[1]^. According to the World Health Organization, schizophrenia affects the health of more than 23 million people. At present, clinical differential diagnosis mainly depends on symptom and disease severity assessment scales, cognitive assessment scales, etc. Therefore, it is urgent to explore objective diagnostic markers and related molecular mechanisms.

Biomarkers are closely related to the course of disease and can be used to monitor changes in disease status ^[2]^, including genetic modification of hormones, proteins, DNA or RNA, and even structural or functional changes visible by imaging techniques. The source of biomarkers for most researchers is blood, which is an important component of human body fluids and is directly connected to various systems and organs of the body. However, blood is strictly regulated by the steady-state mechanism, and it is not easy to accumulate changes in the body before reaching the compensatory critical point of the body. Urine does not directly contact with the body organs, but trace its source is the filtrate of the blood, not regulated by the body ‘s steady-state mechanism, can enrich the blood to produce small changes.

Changes throughout the course of the disease can be found in these small changes and are a treasure trove in the search for disease markers, especially in the early stages of the disease before blood changes, so urine responds to body changes faster and more sensitively than blood.

Proteomics is a quantitative analysis of protein expression in biological samples. It is a powerful tool for identifying new molecular biomarkers and can also provide information on the pathophysiological mechanisms of diseases. Relative quantification by liquid chromatography-tandem mass spectrometry (LC-MS / MS) is a promising protein biomarker identification technique. Proteomics analysis shows that there are considerable pathophysiological changes in depression ^[3,4]^, Alzheimer ‘s disease ^[5]^, Parkinson ‘s disease ^[6]^ and other mental systemic diseases. The purpose of this study is to explore whether urinary proteomics has the potential to identify objective diagnostic markers of schizophrenia, whether it can distinguish between patients and normal people from the perspective of urinary proteomics, and provide clues for clinical diagnosis and other related biomarkers.

## 2 material and methods

### 2.1 Urine collection and ethics statement

A total of 10 first-episode untreated schizophrenic patients and 9 healthy controls were collected from Beijing Anding Hospital. Urine samples were morning urine. This experiment was based on the reuse of discarded samples from the laboratory. The process did not involve any identity information of the patient. We did not affect any treatment of the patient and did not recommend any clinical and auxiliary examinations to the patient. The knowledge generated in this study is in the research stage. All participants signed informed consent and the study was ethically approved by the Ethics Committee of Beijing Anding Hospital, Capital Medical University (No. (2020) Research No. (104)).

### 2.2 urine sample preparation

#### (1) urinary protein extraction

The collected urine samples were temporarily stored in-80 °C refrigerator. Four millilitres of urine sample reheated at 37°C was centrifuged at 4°C and 12,000g for 20min to remove cell and cell debris and retain the supernatants. Next, 20 mM dithiothreitol (DTT) was added and heated at 99.2°C for 10 min to cool to room temperature. Then 50 mM iodoacetamide (IAA) was added and mixed at room temperature for 40 min in the dark. After the reaction finished, the supernatants were placed into new Eppendorf tubes and mixed with four volumes of ethanol at –20°C overnight to precipitate protein. Mixed liquid was centrifuged at 4°C and 10,000g for 30min, and the precipitate was retained and dried. Proper lysis buffer (8 mol/L urea, 2 mol/L thiourea, 50 mmol/L Tris, and 25 mmol/L DTT) was added to redissolve the protein precipitate and was mixed at 4°C for 2h. The supernatants were retained after centrifugation at 4°C and 12,000g for 30min, and each sample’s protein concentration was measured by the Bradford method.

#### (2) urinary protein tryptic digestion

The FASP method^[7]^ was used to digest urine protein with trypsin (Trypsin Gold, Mass Spec Grade, Promega, Fitchburg, Wisconsin, USA). One hundred micrograms of urine protein was added in the membrane of a 10KD ultrafiltration tube (Pall, Port Washington, NY) and washed twice with UA (8 M urea, 0.1 mol/L Tris–HCl, pH 8.5) and NH4HCO3 (25 mmol/L) at 14,000 g for 40 min at 18 °C. Then, the urine protein was resuspended in 25mmol/L NH4HCO3 and digested by trypsin (enzyme to protein ratio of 1:50) at 37 °C for 14 h– 16 h. We obtained the peptides after two centrifugations at 4°C and 12,000g for 30min. These peptides were desalted using Oasis HLB cartridges (Waters, Milford, MA) and then dried by Speed Vac (Thermo Fisher Scientific, Bremen, Germany). Then, the peptides were stored at –80°C.

#### (3) Peptide fractionation

We applied a method to improve the ability to identify low–abundance peptides. The digested peptides were redissolved in 0.1% formic acid and quantified to 0.5 μg/μL which was measured by the BCA method (Thermo Scientific). A pooled sample (73.5 μg, 3.5 μg of each sample) from 21 samples was loaded onto an equilibrated, high–pH, reversed–phase fractionation spin column (84868, Thermo Scientific) to generate the spectral library for subsequent analysis. Two fractions were collected by pooled sample and water. The remaining eight different fractions were collected by a step gradient of increasing acetonitrile concentrations (5.0, 7.5, 10.0, 12.5, 15.0, 17.5, 20.0 and 50.0% acetonitrile) in a volatile high–pH elution solution. Then the ten fractions were dried by Speed Vac (Thermo Scientific, Bremen, Germany) and resuspended in 20 µl of 0.1% formic acid for LC–MS/MS analysis.

### 2.3 LC-MS/MS analysis

The iRT reagent (Biognosys, Switzerland) was used to calibrate the retention time of the extracted peptide peaks and it was added at a ratio of 1:10 v/v to all of the peptide samples. For analysis, 1 μg of each peptide sample was loaded onto a trap column (75 µm × 2 cm, 3 µm, C18, 100 Å) and separated on a reverse–phase C18 column (50 µm × 15 cm, 2 µm, C18, 100 Å) using the EASY–nLC 1200 HPLC system (Thermo Fisher Scientific, Waltham, MA). The elution gradient was 4%–35% buffer B (0.1% formic acid in 80% acetonitrile, flow rate, 0.4 μL/min) for 90 min. Eluted peptides were analysed by using an Orbitrap Fusion Lumos Tribrid Mass Spectrometer (Thermo Fisher Scientific, Waltham, MA, USA).

To generate the spectral library, the ten fractions from the reversed–phase fractionation spin column were analysed by data–dependent (DDA) MS/MS acquisition mode with a resolution of 120,000 in full scan mode and 30,000 in MS/MS mode. The full scan was performed in the Orbitrap from 350–1,500 m/z, and the high–energy collisional dissociation (HCD) energy was set to 30%. The mass spectrometric parameters included the following: the automatic gain control (AGC) target was set to 4e5; the maximum injection time was 45 ms. nineteen single peptide samples were analysed by data–independent (DIA) MS/MS acquisition mode, and the variable isolation window of the DIA method with 36 windows was applied to DIA acquisition. The mass parameters contained the following: the fall scan was obtained from 350 to 1,500 m/z with a resolution of 60,000, and the DIA scan was obtained from 200 to 2,000 m/z with a resolution of 30,000; the HCD energy was set to 32%; the AGC target was set to 1e6 and the maximum injection time was 100 ms.

### 2.4 data analysis

Ten raw files of DDA data collected from the mass spectrometer were searched by Proteome Discoverer (version: 2.1, Thermo Scientific, USA) against the Swiss–Prot Home Sapiens Human database, which also includes the iRT sequence. The retrieval arguments contained: tryptic digestion; two missed trypsin cleavage sites were allowed; the carbamidomethyl of cysteine was set as a fixed modification; the oxidation of methionine was set as a variable modification; the fragment ion mass tolerance was 0.02 Da; and the parent ion mass tolerance was set to 10 ppm. The false discovery rate (FDR) of proteins and peptides was less than 1%.

Nineteen single DIA raw files and ten DDA raw files were generated in Spectronaut™ Pulsar X (Biognosys, Switzerland) software to build a database. Then single DIA raw files were analysed with this database by using the Spectronaut, and the results were filtered depending on having Q–value of less than 0.01 (corresponding to an FDR of 1%). Proteins recognized by more than two unique peptides were retained. The peptide intensity was calculated by summing the peak areas of the respective fragment ions from MS2 and the protein intensity was calculated by summing the intensities of each peptide.

### 2.5 statistical analysis and functional annotation

The differential protein selection parameters were as follows: fold change≥2 or ≤0.5, p values<0.01. Differential protein functional annotation were generated by DAVID (https://david.ncifcrf.gov), and the functions of differential proteins were searched in the reported literature based on the PubMed database (https://pubmed.ncbi.nlm.nih.gov). The mass spectrometry proteomics data have been deposited to the ProteomeXchange Consortium (http://proteomecentral.proteomexchange.org) via the iProX partner repository^[8,9]^with the dataset identifier PXD039108.

## 3 results and discussion

### 3.1 urine proteome analysis of schizophrenia

For the collected two groups of 19 urine samples, after protein extraction and membrane tryptic digestion of urinary protein, LC-MS / MS label-free quantification and DIA data acquisition mode were performed for mass spectrometry analysis. A total of 1945 proteins (specific peptides ≥ 2, FDR < 1 %) were identified. The samples of schizophrenia patients and healthy group were compared. The screening conditions were FC ≥ 2 & ≤ 0.5, P < 0.01. A total of 35 differentially expressed proteins were screened, and ROC curve analysis was performed on each protein. It is worth noting that the aromatic amino acid decarboxylase related to the pathogenesis of schizophrenia can distinguish all 10 patients and 9 normal people with 100 % accuracy in this study. The AUC value of 17 proteins is greater than or equal to 0.9. The specific information of differential proteins is shown in Table 1.

**Table 1.** Differential protein information identified in schizophrenia group compared with healthy group

| Uniprot ID | Protein name | Fold change | P-values | AUC value | Related to schizophrenia |
| --- | --- | --- | --- | --- | --- |
| P20711 | Aromatic-L-amino-acid decarboxylase | 0.35 | 1.16E-03 | 1 | [10] |
| P98164 | Low-density lipoprotein receptor-related protein 2 | 0.5 | 7.43E-04 | 0.9778 | [19] |
| P24666 | Low molecular weight phosphotyrosine protein phosphatase | 2.34 | 7.46E-05 | 0.9667 |  |
| P62328 | Thymosin beta-4 | 4.14 | 2.52E-03 | 0.9556 | [17] |
| Q9Y4K1 | Beta/gamma crystallin domain-containing protein 1 | 3.13 | 1.29E-03 | 0.9556 |  |
| P07737 | Profilin-1 | 2.96 | 2.09E-03 | 0.9222 | [22] |
| P51654 | Glypican-3 | 0.37 | 3.73E-03 | 0.9222 | [14] |
| P13727 | Bone marrow proteoglycan | 0.35 | 3.38E-03 | 0.9222 |  |
| P63241 | Eukaryotic translation initiation factor 5A-1 | 3.27 | 3.93E-03 | 0.9111 |  |
| P10606 | Cytochrome c oxidase subunit 5B | 2.19 | 1.32E-03 | 0.9111 |  |
| P27482 | Calmodulin-like protein 3 | 6.66 | 3.65E-03 | 0.911 | [11] |
| P14780 | Matrix metalloproteinase-9 | 6.58 | 4.55E-03 | 0.911 | [12] |
| P25815 | Protein S100-P | 2.8 | 5.21E-03 | 0.9 |  |
| P54727 | UV excision repair protein RAD23 homolog B | 2.45 | 2.30E-03 | 0.9 |  |
| P29323 | Ephrin type-B receptor 2 | 0.47 | 3.60E-04 | 0.9 |  |
| Q93088 | Betaine--homocysteine S-methyltransferase 1 | 0.4 | 1.42E-03 | 0.9 |  |
| O75882 | Attractin | 0.39 | 1.07E-03 | 0.9 |  |
| Q14CN2 | Calcium-activated chloride channel regulator 4 | 4.92 | 2.25E-03 | 0.8889 |  |
| Q09666 | Neuroblast differentiation-associated protein AHNAK | 4.8 | 7.52E-03 | 0.8889 |  |
| Q06278 | Aldehyde oxidase | 3.91 | 6.53E-03 | 0.8889 | [20] |
| P08473 | Neprilysin | 0.49 | 1.82E-03 | 0.8889 | [18] |
| Q96DA0 | Zymogen granule protein 16 homolog B | 0.27 | 1.03E-03 | 0.8889 |  |
| P16422 | Epithelial cell adhesion molecule | 3.25 | 3.88E-03 | 0.8778 |  |
| O15231 | Zinc finger protein 185 | 2.9 | 3.30E-03 | 0.8778 |  |
| P46108 | Adapter molecule crk | 2.59 | 6.29E-03 | 0.8778 |  |
| P54802 | Alpha-N-acetylglucosaminidase | 0.45 | 8.12E-03 | 0.8667 | [23] |
| P05787 | Keratin, type II cytoskeletal 8 | 3.16 | 8.13E-03 | 0.8444 |  |
| P19320 | Vascular cell adhesion protein 1 | 2.68 | 7.23E-03 | 0.8444 | [13] |
| O95171 | Sciellin | 2.3 | 7.03E-03 | 0.8444 |  |
| O75487 | Glypican-4 | 0.41 | 4.06E-03 | 0.8444 | [15] |
| Q16610 | Extracellular matrix protein 1 | 5.53 | 4.55E-03 | 0.844 | [16] |
| P46940 | Ras GTPase-activating-like protein IQGAP1 | 3.11 | 4.83E-03 | 0.8333 |  |
| O14950 | Myosin regulatory light chain 12B | 2.96 | 9.36E-03 | 0.8222 |  |
| Q6UWP8 | Suprabasin | 2.86 | 8.94E-03 | 0.8222 |  |
| P02656 | Apolipoprotein C-III | 2.6 | 5.88E-03 | 0.8111 | [21] |

Among the 35 differential proteins identified, protein function analysis and literature search revealed that aromatic-L-amino-acid decarboxylase (AADC) was reported to be related to the pathogenesis of schizophrenia ^[10]^. Calmodulin-like protein 3 ^[11]^, matrix metalloproteinase-9 ^[12]^ and Vascular cell adhesion protein 1 (VCAM1) have been directly reported to be related to schizophrenia. The expression of VACM1 in schizophrenia is up-regulated in the literature ^[13]^, consistent with our results. Many of these proteins have important regulatory roles in neural development and synaptic plasticity, such as Glypican-3 ^[14]^ and Glypican-4 ^[15]^, extracellular matrix protein 1 ^[16]^ ; some proteins have also been reported to be related to other neurological diseases. For example, thymosin beta-4 has been reported to be used in the treatment of other neurological diseases ^[17]^. Neprilysin (NEP) is an important neuropeptidase and amyloid degrading enzyme, which is reported to be a therapeutic target for Alzheimer ‘s disease ^[18]^. There are also some proteins associated with drug metabolism for the treatment of schizophrenia, such as low-density lipoprotein receptor-related protein 2 ^[19]^, aldehyde oxidase ^[20]^ and apolipoprotein C-III ^[21]^ ; in addition, in the study of prenatal stress mouse model, it was found that the expression of profilin-1 in the hippocampus of prenatal stress mice was changed ^[22]^, and prenatal stress can affect the development of the brain and increase the risk of mental illness in the future. In patients with schizophrenia and depression, the phenomenon of abnormal hippocampus has been clarified, which indicates that profibrin-1 is involved in the pathogenesis of schizophrenia. At the same time, in a case report, it was mentioned that the activity of Alpha-N-acetylglucosaminidase (NAGLU) was basically undetectable when a patient developed mental illness ^[23]^. This may suggest that we use NAGLU as a potential biomarker for further research.

### 3.2 Results of random generation analysis

In order to determine the possibility of random generation of identified differentially expressed proteins, we randomly grouped the total proteins identified in 19 samples of two groups, and applied the same criteria for screening differentially expressed proteins : FC ≥ 2 & ≤ 0.5, P < 0.01. After 92378 random groupings, the average number of differentially expressed proteins was 3.01, and the proportion of randomly identified proteins was 8.60 %, indicating that at least 91.40 % of the differentially expressed proteins were not due to randomness. The results of the random grouping test are shown in Table 2.The probability that the 35 differential proteins we screened can be randomly generated is low, and it is inferred that these differential proteins are indeed related to schizophrenia.

**Table 2.** Results of random generation analysis

| Screening criteria | Total number of random combinations | Average number of proteins with false random combinations | Ratio<br>(average numbers of proteins with false random combinations/number of correctly identified differential proteins) |
| --- | --- | --- | --- |
| FC≥2 or ≤0.5<br>P < 0.01 | 92378 | 3.01 | 0.086 |

### 3.3 Functional annotation

Functional annotation analysis of differential proteins was performed using DAVID to analyze the biological processes, cellular components, and molecular functions associated with differential proteins (Figure 2). From the biological process part of the map, it can be seen that these differentially expressed proteins are mainly involved in cell migration, ephrin receptor signaling pathway, regulation of signal transduction, tumor necrosis factor-mediated signaling pathway, regulation of T cell migration, regulation of protein localization to membrane, cellular response to UV-A, β-amyloid clearance, negative regulation of peptidase activity, positive regulation of long-term synaptic potentiation(LTP) and lipid metabolism process. Among them, pathway-related proteins that regulate the transmission of nerve impulses, such as ephrin receptor signaling pathway and positive regulation of LTP, have been reported to be related to the occurrence of certain brain diseases ^[24].^ From the cellular component part of the diagram, it can also be seen that these differential proteins are mainly derived from the cell muscle fiber membrane, axons, etc., indicating that these differential proteins are significantly related to the transmission of nerve impulses.

## 4 Conclusion

There are significant differences in urine proteome between schizophrenia and healthy people. A number of potential high-quality biomarkers of schizophrenia have been found, which also provide clues for the study of pathogenesis and the discovery of targets.

---

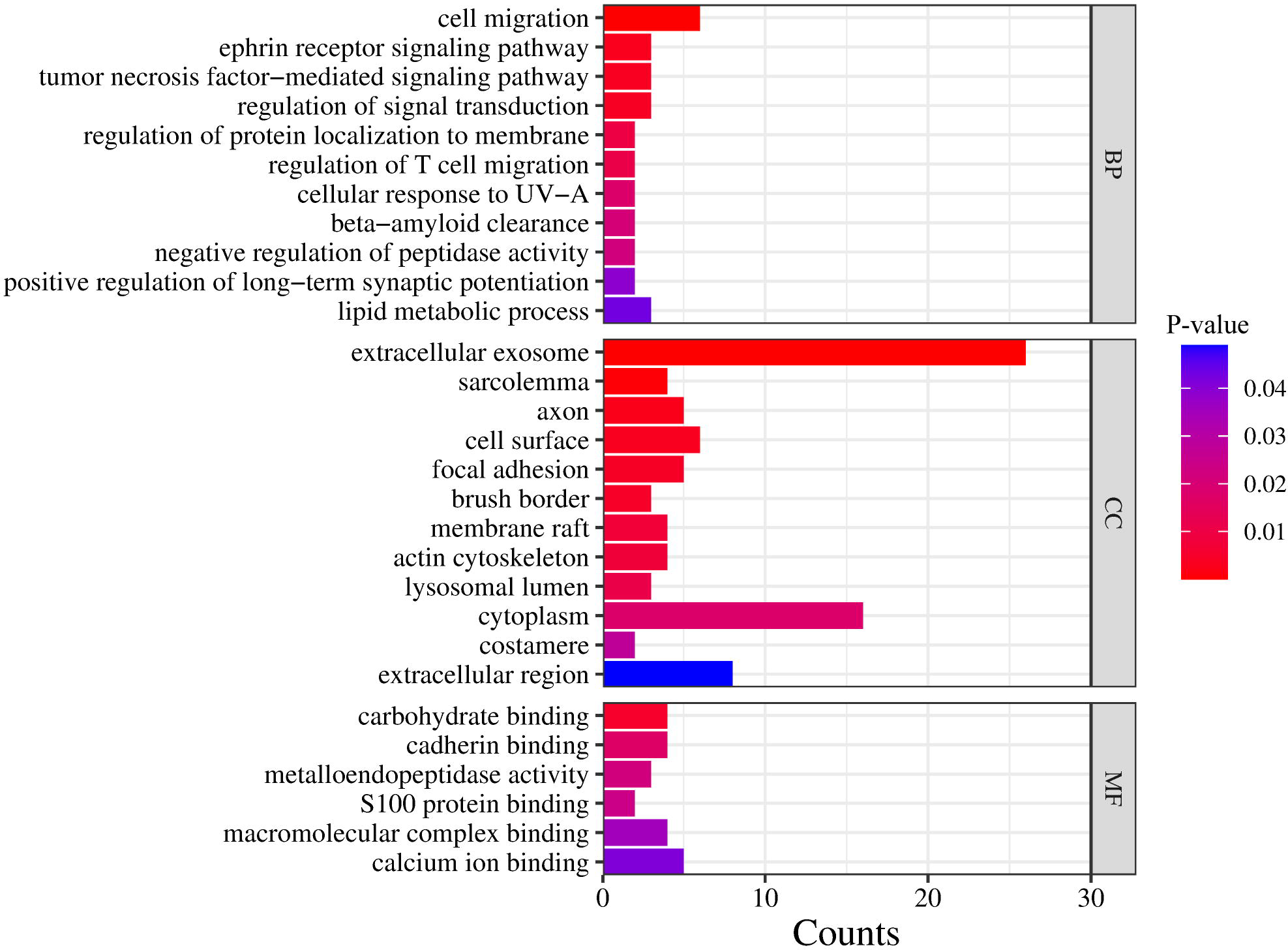

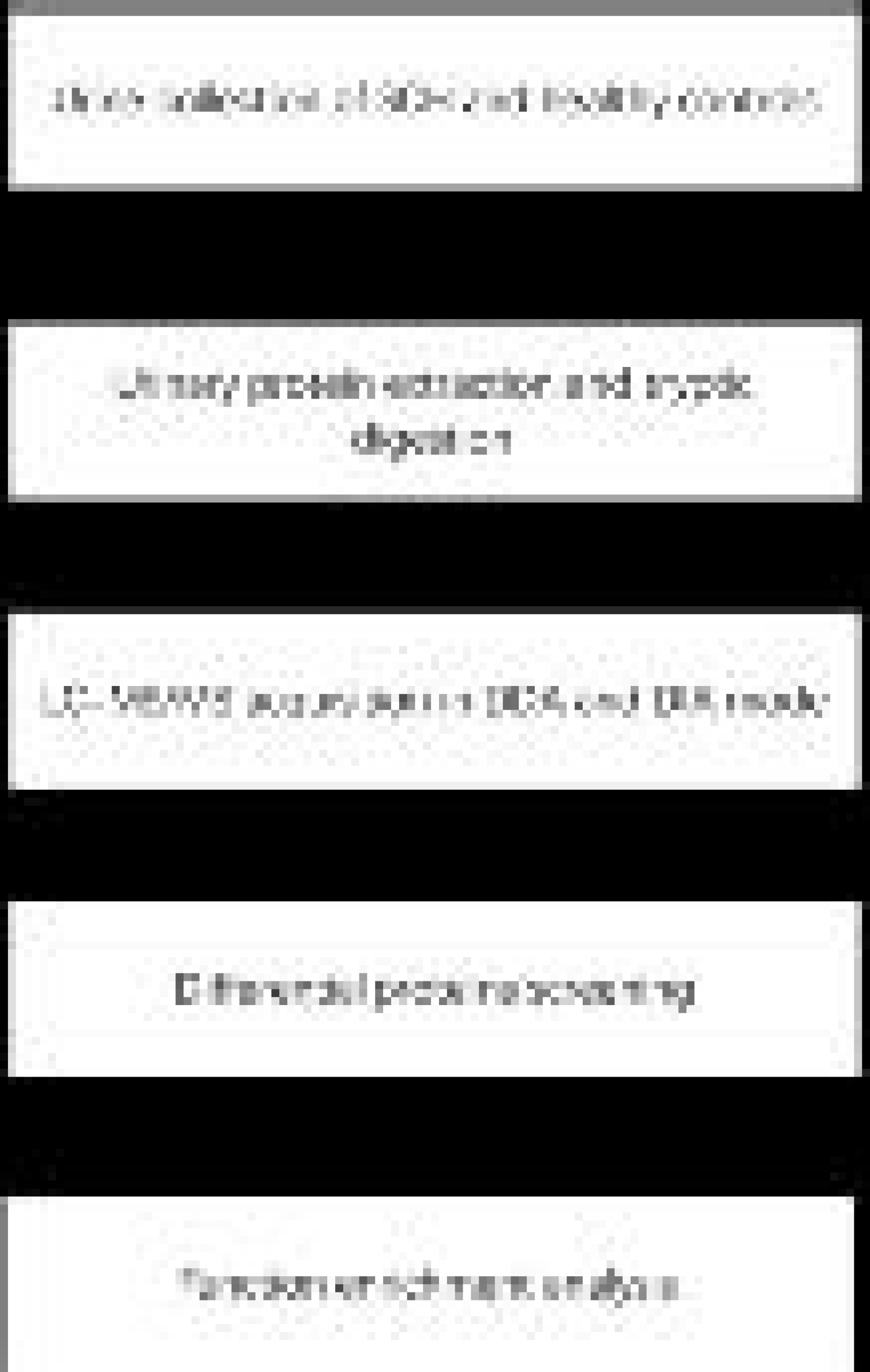

